# New human chromosomal safe harbor sites for genome engineering with CRISPR/Cas9, TAL effector and homing endonucleases

**DOI:** 10.1101/396390

**Authors:** Stefan Pellenz, Michael Phelps, Weiliang Tang, Blake T. Hovde, Ryan B. Sinit, Wenqing Fu, Hui Li, Eleanor Chen, Raymond J. Monnat

## Abstract

Safe Harbor Sites (SHS) are genomic locations where new genes or genetic elements can be introduced without disrupting the expression or regulation of adjacent genes. We have identified 35 potential new human SHS in order to substantially expand SHS options beyond the three widely used canonical human SHS, *AAVS1, CCR5* and *hROSA26*. All 35 potential new human SHS and the three canonical sites were assessed for SHS potential using 9 different criteria weighted to emphasize safety that were broader and more genomics-based than previous efforts to assess SHS potential. We then systematically compared and rank-ordered our 35 new sites and the widely used human *AAVS1, hROSA26* and *CCR5* sites, then experimentally validated a subset of the highly ranked new SHS together versus the canonical *AAVS1* site. These characterizations included *in vitro* and *in vivo* cleavage-sensitivity tests; the assessment of population-level sequence variants that might confound SHS targeting or use for genome engineering; homology–dependent and –independent, SHS-targeted transgene integration in different human cell lines; and comparative transgene integration efficiencies at two new SHS versus the canonical *AAVS1* site. Stable expression and function of new SHS-integrated transgenes were demonstrated for transgene-encoded fluorescent proteins, selection cassettes and Cas9 variants including a transcription transactivator protein that were shown to drive large deletions in a *PAX3/FOXO1* fusion oncogene and induce expression of the *MYF5* gene that is normally silent in human rhabdomyosarcoma cells. We also developed a SHS genome engineering ‘toolkit’ to enable facile use of the most extensively characterized of our new human SHS located on chromosome 4p. We anticipate our newly identified human SHS, located on 16 chromosomes including both arms of the human X chromosome, will be useful in enabling a wide range of basic and more clinically-oriented human gene editing and engineering.

## Introduction

Safe harbor sites (SHS) are genomic loci where genes or other genetic elements can be safely inserted and expressed. These SHS are critical for effective human disease gene therapies; for investigating gene structure, function and regulation; and for cell marking and tracking. The most widely used human SHS were identified by serendipity (e.g., the *AAVS1* adeno-associated virus insertion site on chromosome 19^1,2^); by homology with useful SHS in other species (e.g., the human homolog of the murine *Rosa26* locus^3^); and most recently by recognition of the dispensability of a subset of human genes in most or all individuals (e.g., the *CCR5* chemokine receptor gene, that when deleted confers resistance to HIV infection^4–6^).

In order to more systematically identify and expand the number of useful human SHS, we first searched for target sites in the human genome predicted to be efficiently cleaved by the canonical genome engineering homing/meganuclease mCreI^7–9^. We reasoned that any potential SHS identified by the presence of a high quality mCreI site would also contain one or more adjacent cleavage sites for Cas9 and TALEN genome engineering nucleases that have less stringent targeting requirements. This initial precise anchoring of potential new safe harbor sites in the human genome would, in turn, facilitate better assessment of site safety, potential functional competence and the presence of confounding sequence variations. Our aim was to identify new human SHS that could be targeted by any of these three nuclease types, in a wide range of human cells, to broadly enable a wide range of basic science as well as clinical applications.

We report here the identification of 35 potential new human SHS, located on 16 different human chromosomes and 23 chromosome arms including both arms of the human X chromosome. These 35 new SHS and the three canonical human SHS (*AAVS1*, the human *ROSA26* locus and *CCR5)* were assessed and rank-ordered for safety and potential utility using a comprehensive scoring system that included 8 different genomic criteria in addition to uniqueness. Several high-ranking potential new SHS were experimentally validated by PCR amplification, mCreI cleavage sensitivity and DNA sequencing, together with a demonstration of efficient editing and transgene insertion mediated by Cas9, TALEN and mCreI nucleases. SHS-specific transgene insertion by both homology-mediated as well as cleavage-dependent, likely homology-independent mechanisms was demonstrated. The most extensively characterized of these new SHS, the high-ranking SHS231 located on the proximal long arm of chromosome 4, was also shown to be functionally competent for recombinase/integrase-mediated editing. Selectable, scorable and fluorescent/functional protein-encoding SHS231 transgenes were shown to be stably expressed when compared with the same transgenes inserted into the canonical *AAVS1* site in a number of different human cell lines. The SHS231 engineering toolkit will allow others to make rapid use of this enhanced chromosome 4 SHS for both basic and clinically-oriented genome engineering applications.

## Materials and Methods

### Cell lines/cell culture

Human 293T cells or derivatives and four human rhabdomyosarcoma (RMS) cell lines derived from unrelated patients were used for experiments. All five lines were cultured in D-MEM medium supplemented with 10% (v/v) fetal bovine serum (Hyclone, GE Healthcare/ Biosciences, Pittsburgh, PA), 2 mM L-glutamine and antibiotics (1% Pen-Strep, Gibco, Thermo Fisher Scientific, Waltham, MA) in a 5% CO_2_ humidified 37°C incubator. Human 293T-REX cells, a derivative of the parent 293T cell line (ATCC cell line CRL-3216), were grown in accordance with the supplier’s instructions (Invitrogen/Thermo Fisher, Waltham, MA). The human RMS cancer cell lines RD, Rh5, Rh30 and SMSCTR have been described previously^10^, and were obtained the laboratories of Dr. Corinne Linardic (Duke University School of Medicine, Durham, NC) and Dr. Charles Keller (Children’s Cancer Therapy Development Institute, Beaverton, OR). Cells were tested periodically for *Mycoplasma* infection and authentication was done by DNA fingerprinting (the RMS lines were verified by the Dana Farber Cancer Institute Molecular Diagnostic Laboratory by short tandem repeat profiling).

### SHS identification and experimental validation

In order to identify potential new human SHS, we first searched the human genome for high quality matches to the target sequence of the canonical homing endonuclease mCreI. We reasoned that a SHS identified by a highly cleavage-sensitive mCreI target site or variant would also contain one or more adjacent cleavage sites for Cas9 and TALEN-based nucleases that have less stringent targeting requirements. The well-defined mCreI site would also anchor the search of adjacent chromosomal DNA to assess and rank-order SHS suitability based on criteria for site safety, functional competence and the presence of potentially confounding sequence variations. This search was initiated by using detailed information on the cleavage specificity of mCreI that quantified the contribution of each basepair in the mCreI target site sequence. This position weight matrix was used to construct a list of 128 target site sequence variants predicted to be cleaved with ≥90% of the efficiency of the native mCreI site^11–16^ (Fig. 1A and 1B). These 128 mCreI target site variants were FASTA-formatted and uploaded to the NCBI BLAST search engine (http://blast.ncbi.nlm.nih.gov/) in order to identify target site matches in the human genome (GRCh37/hg19) using the following BLAST parameters: optimize for ‘Highly similar sequences (megablast)’; max target seqs = 50; short queries: ‘adjust for short sequences’; expect threshold = 1; word size = 7; match/mismatch: 4, −5; and gap cost: existence=12/extension= 8. All resulting genomic target site matches of ≥95% identity (19/20 or 20/20 bp matches versus the canonical mCreI target site) were subsequently evaluated as potential new safe harbor sites.

**Figure 1.**
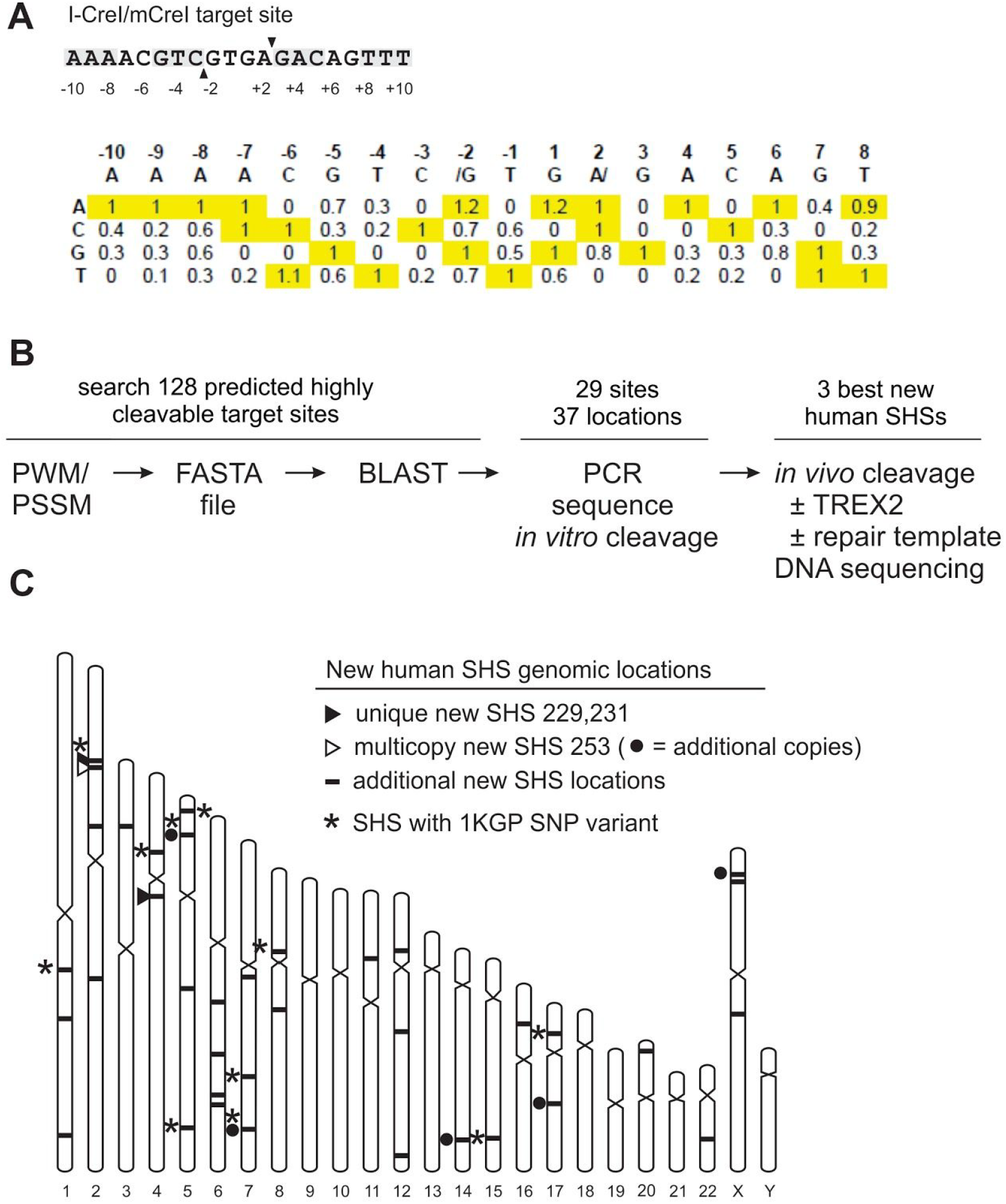
Identification and mapping of new human safe harbor sites (SHS). **(A)** The canonical mCreI homing endonuclease cleavage site is shown top with twofold symmetric basepair positions shaded. The matrix below summarizes the functional consequences of basepair insertions across the mCreI target site where a value of 1 = native site cleavage efficiency and values < 0.3 indicate cleavage resistance. Basepairs highlighted in yellow indicate either the canonical basepair at that position, or a highly cleavable basepair substitution. **(B)** Workflow for identifying highly cleavage-sensitive mCreI target sites in the human genome sequence. **(C)** Physical confirmation and functional verification of two new unique SHS located on chromosomes 2p (SHS229) and 4q (SHS231). A third highly ranked SHS (SHS253) was identified at 6 locations on the short arms of chromosomes 2, 5 and X and the long arms of chromosomes 7, 14 and 17. Asterisks (*) indicate sites where basepair variants have been identified in the mCreI target site in human population genetic data (see text for additional detail).

Potential new human SHS identified by BLAST search and the canonical human SHS *AAVS1, HsROSA26* and *CCR5* were then evaluated for SHS potential by 8 criteria in addition to site uniqueness that assessed site safety, accessibility and functional criteria (Fig. 1C; Table 1 and 2). These criteria were based on several less extensive lists of criteria (e.g., proximity to known genes or regulatory elements, see, *e.g.*, Sadelain et al 2012^17^), and made use of contemporary genomic data, *e.g.*, ENCODE Consortium project results^18^. All SHS candidates including the three canonical human SHS were evaluated as follows: sites were first searched 300 kb up-and downstream in the UCSC Genome Browser in order to identify genes or RNAs, especially any already related to cancer; proximity to any transcriptionally active region regardless of annotation; the presence of replication origins or ultra-conserved elements; location in open chromatin as assessed by nuclease sensitivity; and whether the SHS was located in a region of copy number variation^19,20^ (CNV; http://genome.ucsc.edu/). We next used 1000 Genomes Project (1KGP) data (http://www.ncbi.nlm.nih.gov/variation/tools/1000genomes/) to identify basepair-level population genetic variation within all of the mCreI-anchored SHS sites ^21^ (Table S2). This approach was used to provide an estimate of the fraction of SHS that would be directly accessible in individuals by mCreI (and, by extension, other genome engineering nucleases). New SHS that differed from the canonical mCreI site at 1 or more basepair positions were further assessed using the mCreI position weight matrix (PWM) developed from single base-pair profiling experiments^14,16^ (Fig. 1B) to predict cleavage sensitivity.

**Table 1.** Criteria for identification and assessment of new human Safe Harbor Sites (SHS)

|  | SHS criterion | UCSC browser track source |
| --- | --- | --- |
| **safety** | 1. > 300kb from any cancer-related gene on allOnco list | genes and gene predictions: UCSC Genes |
|  | 2. > 300kb from any miRNA/other functional small RNA | genes and gene predictions: sno/miRNA |
|  | 3. > 50kb from any 5' gene end | genes and gene predictions: RefSeq Genes |
| **functional silence** | 4. > 50kb away from any replication origin | regulation: UW Repli-seq: Peaks |
|  | 5. > 50kb away from any ultraconserved element | regulation: VISTA Enhancers |
| | 6. low transcriptional activity (no mRNA ± 25kb) | mRNA and EST: Human mRNAs |
| **consistent/**<br>**accessible/**<br>**unique** | 7. not in copy number variable region | repeats: Segmental Dups |
| | 8. in open chromatin (DHS signal ± 1kb) | regulation: ENC DNase/FAIRE: Uniform DNaseI HS |
|  | unique (1 copy in human genome) | BLAST search output |

**Table 2.**
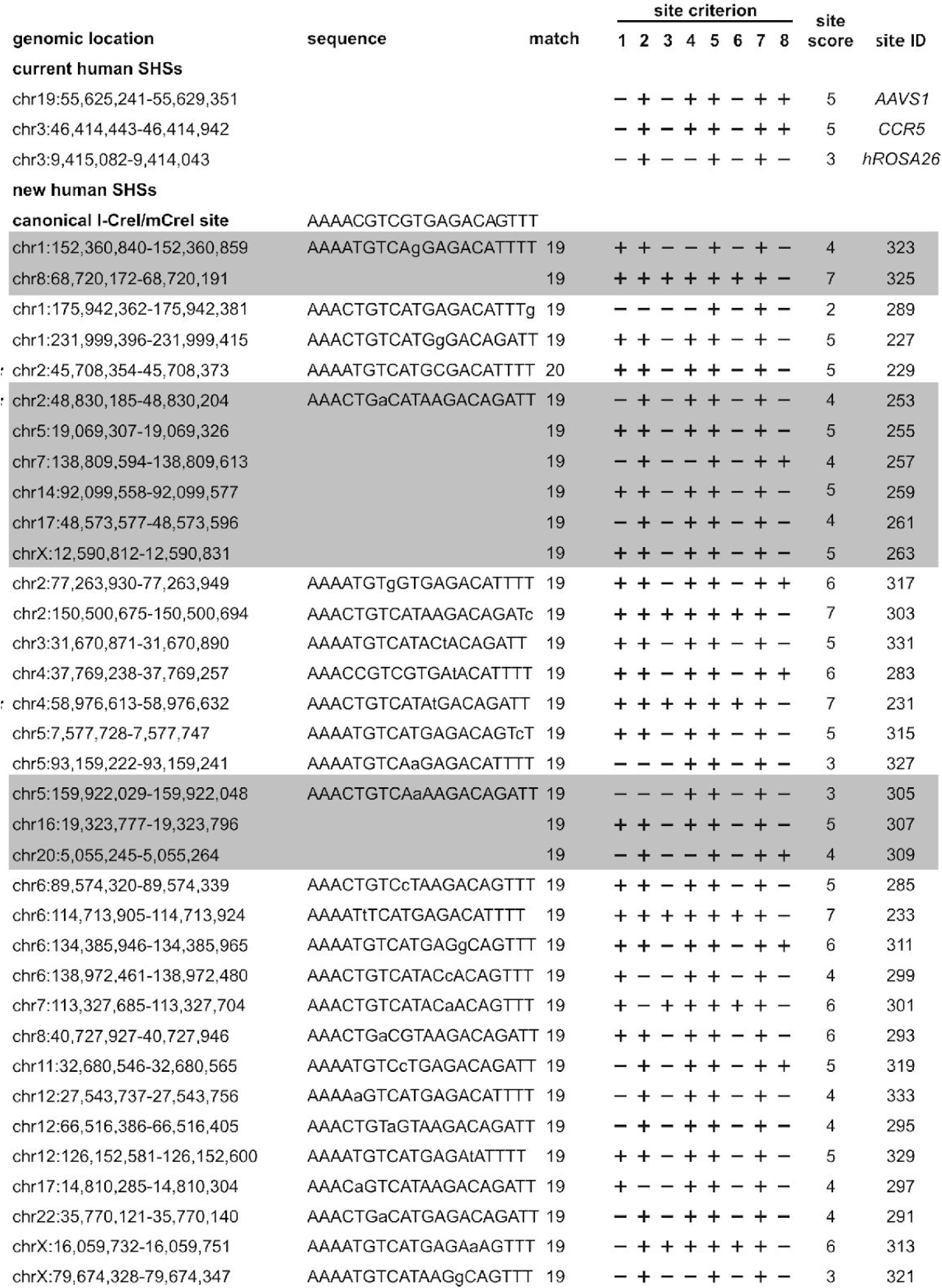
Safe harbor site scoring

Potential new SHS identified and assessed by the above criteria were then rank-ordered and experimentally validated by PCR amplification and mCreI *in vitro* cleavage analyses. Site-specific primer pairs were designed using CLC Workbench Primer Design Tool (http://www.clcbio.com; CLC Bio, Boston, MA) to generate ∼300-400 bp PCR products containing the mCreI target site (Table S1). Genomic DNA purified from human 293T cells using a Wizard Genomic DNA Purification Kit (Promega, Madison, WI) was used as the template for SHS amplifications (Table S1). SHS amplification reactions were performed in 25 μL of 1X Thermo polymerase buffer containing all four dNTPs at 200 µM, 150 ng of genomic DNA and 400 nM of each primer with 1.25 units of Taq polymerase (New England Biolabs; NEB, Ipswich, MA). Amplifications were performed using a 1 min 95°C denaturation step followed by 30 cycles of 30 sec at 95°C; 30 sec at 50°C; and 30 sec at 68°C followed by 5 min at 68°C. Alternatively, a subset of SHS was amplified in 25 μL reactions that contained 12.5 µL PrimeStar Max DNA polymerase premix (Takara, Mountain View, CA), 50 ng of purified genomic DNA and 240 nM final concentration for each amplification primer. Amplifications were performed using 35 cycles of 10 sec at 98°C; 15 sec at 50°C and 3 min at 72°C. SHS-specific PCR products were gel-purified using a QIAquick Gel Extraction Kit (Qiagen, Hilden, Germany), quantified by spectrophotometry, then digested with purified mCreI protein in 15 μL reactions containing 15 fmol DNA substrate and 0, 15 or 150 fmol of purified mCreI protein^8,16^ in 170 mM KCl, 10 mM MgCl_2_ and 20 mM Tris pH 9.0. Digestions were performed at 37°C for 1 hr, then stopped by adding 3 μL (1:6) of 6x stop buffer (60 mM Tris.HCl pH 7.4, 3% SDS, 30% glycerol, 150 mM EDTA) prior to electrophoresis through a 1% agarose gel run in TAE buffer (40 mM Tris, 20 mM acetic acid, 1 mM EDTA). Substrate and cleavage product bands were identified following gel electrophoresis by ethidium bromide staining, digital image capture and band intensity quantification using ImageJ (http://imagej.nih.gov/ij/). A comparably-sized PCR product containing the native mCreI target site was included in experiments as a positive digestion control. A subset of newly identified SHS were also sequence-verified from PCR products using SHS-specific primers by capillary sequencing (Table S1; Genewiz, South Plainfield, NJ). Sequenced reads were aligned to genomic sequence using CLC Workbench Alignment tool (CLC Bio, Boston, MA).

We verified the *in vivo* cleavage sensitivity of several potential SHS by co-expressing the mCreI homing endonuclease together with the TREX2 3′ to 5′ repair exonuclease in 293T cells. The inclusion of TREX2 allows a more accurate measure of the fraction of sites cleaved *in vivo* by promoting NHEJ-mediated mutagenic repair following site cleavage^22^ (Fig. S1). The expression vector used in these experiments was constructed in a pRRL-based lentiviral vector backbone that encoded the open reading frames for mCreI, the TREX2 exonuclease and mCherry fluorescent protein in a single translational unit separated by self-cleaving T2A peptides^25^ (Fig. S1). Target site cleavage was estimated by amplifying sites from transfected cells, then determining the fraction of PCR products that were mCreI cleavage-resistant and mutant. We extensively analyzed three new SHS in this way: SHS231, a unique chromosome 4 site with the highest SHS score; SHS229, a chromosome 2 SHS with perfect nucleotide sequence identity to a member of our 20 bp site query library; and SHS253, the chromosome 2-specific member of the small family of 6 identical target sites represented once each on 6 different chromosomes (chromosomes 2, 5, 7,14,17 and X; Fig. 1C, Table 2).

A modified calcium phosphate (CaPO_4_) transfection protocol^23^ was used to introduce a pRRL-based lentiviral expression vector encoding mCreI, TREX2 and mCherry proteins into human 293T cells^24^ (Fig. S1). Cells (2-4 x 10e5/well) were plated in a 6-well plate 24 hr prior to transfection and were ∼70% confluent at the time of transfection. Expression vector plasmid DNA (1.5 µg in 10 µL H_2_O) was mixed with 40 µL of freshly prepared 0.25 M CaCl_2_ and 40 µL of 2x BBS buffer (50 mM BES pH 6.95 (NaOH), 280 mM NaCl, 1.5 mM Na_2_HPO_4_; Boston BioProducts), then incubated at room temperature for 15 min before being added dropwise to wells. Plates were incubated overnight in 3% CO_2_ at 37°C. The medium was changed the following day, and cells were grown for an additional 24 hr in a 5% CO_2_, 37°C humidified incubator. Transfection efficiency was checked by determining the fraction of mCherry-positive cells by flow cytometry: in brief, cells were trypsinized, counted and fixed with formaldehyde (1% v/v final concentration, 10 min at room temperature followed by the addition of 1/20 volume of 2.5 M glycine) prior to flow cytometric analysis of ∼2 x 10e4 cells/transfection on a BD FACS Canto II flow cytometer (BD Biosciences, San Jose, CA). Genomic DNA prepared from co-transfected and control cells was used for PCR amplification and *in vitro* mCreI cleavage analysis of specific SHS as described above.

### Homology-dependent SHS editing by three genome engineering nucleases

The mCreI-I expression vector described above, together with SHS231-specific TALEN and CRISPR/Cas9 expression vectors, were used for SHS editing experiments. The SHS231-specific TALEN protein pair was designed using the TALEN Targeter 2.0 web design engine^26,27^ (https://tale-nt.cac.cornell.edu/node/add/talen). Forward and reverse strand, 20 bp-specific TALEN sequences were inserted into the TALEN expression vector pRKSXX-pCVL-UCOE.7-SFFV-BFP-2A-HA-NLS2.0-TruncTAL (kindly provided by Dr. Andrew Scharenberg, Seattle Children’s Research Institute, Seattle WA), and each TALEN open reading frame was generated by assembling the following repeat variable di-residues (RVDs): left TALEN: NG NG NN NN HD NG NI NH NN NH HD NG NI NI NN NN NI NG NG NI, corresponding to the nucleotide sequence TTGGCTAGGGCTAAGGATTA (chr 4: 58,976,594-58,976,613); and right TALEN: NG NN NG NI NG NH HD NG NG NG HD HD NG HD NG NG NN NG NG NI, corresponding to the nucleotide sequence TGTATGCTTTCCTCTTGTTA^26,28^ (chr 4:58,976,613-58,976,632).

A SHS231-specific CRISPR/Cas9 expression vector was constructed in pX260^29,30^ that contained expression cassettes for the *S. pyogenes* Cas9 nuclease, the CRISPR RNA array, and the tracrRNA. The SHS231 Cas9 target site, 5’-AAAACATTTATATACTGCGTGG-3’, was located 110 bp downstream of the mCreI/TALEN cleavage site, was identified using the CRISPR Design Tools Resource developed by Zhang and colleagues^29,30^ (http://crispr.mit.edu/). A corresponding SHS231-specific Cas9 nickase expression vector was also constructed in pX334, which encoded a Cas9 D10A substitution to confer nickase activity. A guide RNA template sequence, 5′-CTAATCTGGACAAAACATTTATATACTGCG-3′, was inserted into both expression vectors followed by a TGG proto-spacer adjacent (PAM) motif^29,30^.

In order to determine whether SHS cleavage *in vivo* could catalyze homology-directed repair in the presence of a homologous donor template, we co-transfected human 293T cells with a SHS-specific repair template and an expression vector for mCreI, for a TALEN pair, or for Cas9 cleavase/nickase enzymes (Fig. 2, Fig. S1). The template for SHS-specific, homology-dependent repair consisted of 500 bp homology arms that flanked the mCreI target site region and contained a 48 bp insert at the center harboring a canonical *loxP* recombinase site and adjacent, diagnostic restriction endonuclease cleavage sites for PvuI and SacII (Fig. 2). Repair templates were made by overlap extension PCR using oligonucleotide primers to generate PCR products that, when re-amplified, incorporated the 48 bp *loxP* insert at the center of the repair template (Table S1).

**Figure 2.**
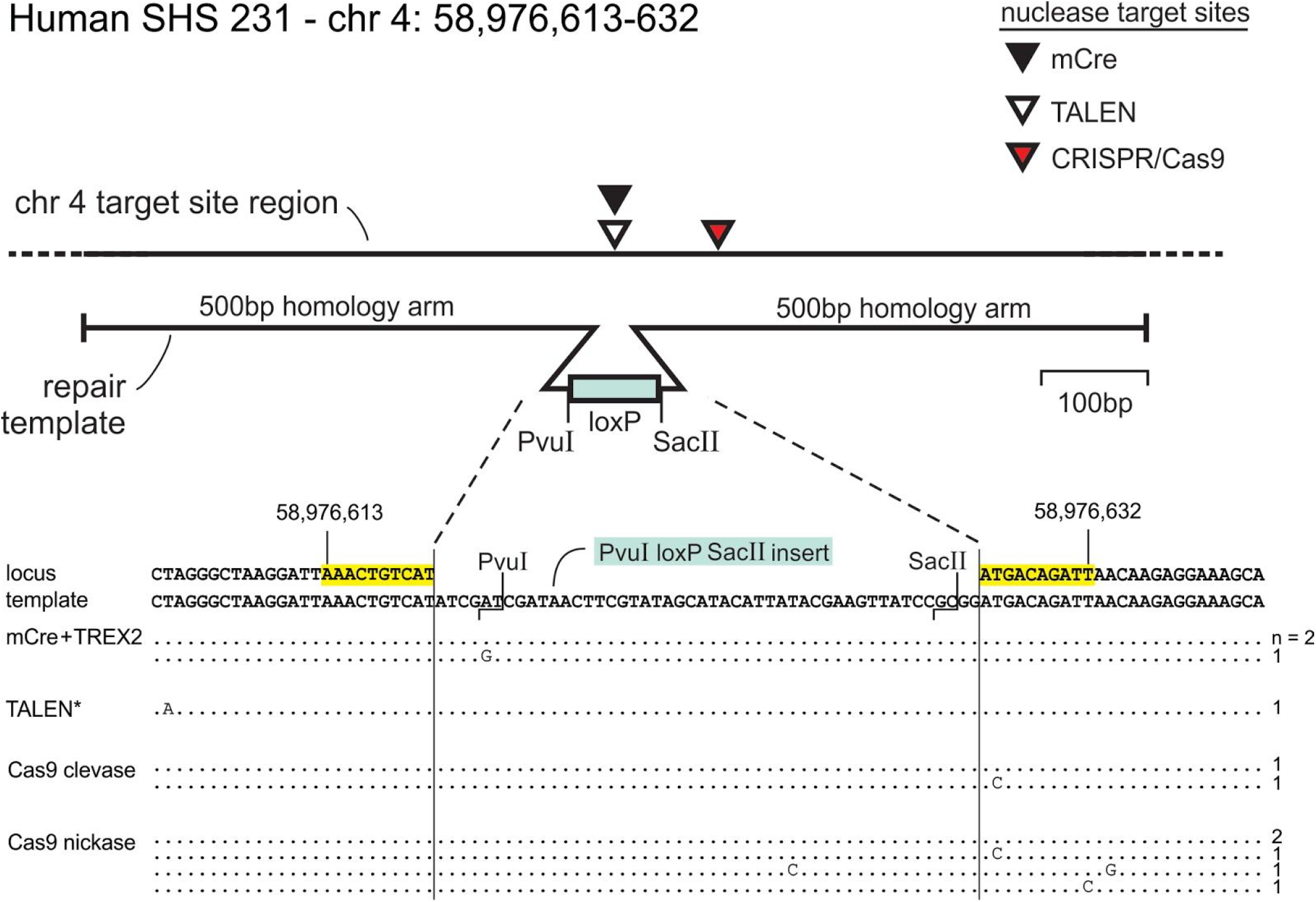
Molecular confirmation of SHS231 homology-dependent editing by three engineering nucleases. The top panel shows the locations of cleavage sites for mCreI, TALEN and CRISPR/Cas9 nucleases centered on the chromosome 4 SHS231 safe harbor site (key shown top right), with the structure of the 1.05 kb repair template shown below. The bottom panel shows independently cloned and sequenced inserts from targeted SHS231 insertions by all 3 nucleases. The mCreI targeting experiments used an expression vector that encoded both mCreI and the TREX2 nuclease (see text), and Cas9 targeting was performed using a common guide RNA and either a Cas9 cleavase or nickase, Numbers to the right of each row indicates the number of independent targeting events that were cloned and sequenced (see text for additional detail).

Calcium phosphate transfection (as described above) was again used to introduce nuclease expression vectors into human 293T cells^24^. Transfection efficiency was checked by determining the fraction of mCherry-positive cells by flow cytometry, as described above. Molecular characterization of SHS editing was performed by PCR amplifying the SHS region of interest from transfected cells, followed by PvuI or SacII restriction digest to confirm targeted integration of the *loxP* cassette (Fig. 2, Fig. S2). PCR products were also cloned into a pGEM-T Easy plasmid vector (Promega, Madison, WI) and transformed into α-Select Chemically Competent Gold Efficiency cells (Bioline, Taunton, MA), followed by plasmid preparation from white (insert-containing) colonies for capillary sequencing using a T7 promoter sequencing primer (Fig. 2). Sequencing results were aligned with the repair template sequence using the CLC Main Workbench software (CLCBio).

### Homology-independent SHS genome editing by Cas9

Homology-independent editing of the SHS231 locus was performed using the protocol above with modified Cas9 and repair template constructs. Dual human U6-driven guide RNAs (gRNA) targeting SHS231 were simultaneously inserted into a custom *S. pyogenes* Cas9-T2A-GFP expression plasmid (pUS2-SH231) using Gibson assembly, as previously described^31^. SHS231-specific gRNAs (SHS231 gRNA1: 5’-GCCTCCCCCATAGTACCAT-3’; SH231 gRNA2: 5’-GATGTGCTCACTGAGTCTGA-3’) were designed to target and cleave both the SHS231 genomic locus and the repair template to promote efficient transgene integration by NHEJ-mediated DNA end joining^32,33^. The transgene cassettes were also flanked by Bxb1 recombinase and ΦC31 *attP* integrase target sites that, once integrated, could be used for high efficiency SHS-specific editing by these recombinase/integrase proteins.

To engineer SHS231 using homology-independent approaches, repair templates (3 µg) and the pUS2-SH231 dual guide-targeting Cas9 expression vector (3 µg) were co-electroporated into three different human rhabdomyosarcoma (RMS) cell lines (Rh5, Rh30, and SMSCTR^10^; 1 x 10e6 cells per transfection) using the 100ul Neon electroporation system (Life Technologies, Carlsbad, CA) according to the manufacturer’s protocol and two, 1150V pulses for 30 ms each. After 2 weeks of selection (puromycin, hygromycin or blasticin, depending on the repair template; see Fig. 1, Table S3), transgene integration was confirmed with PCR amplification of the SHS231 target site (Q5 polymerase, NEB, Ipswich, MA) using a transgene and adjacent genome-anchored primer pair (SHS231 gFwd: GAACCAGAGCCACCCAGTTG, and Bxb1 rev; GTTTGTACCGTACACCACTGAGAC).

### Stable gene expression from SHS231 transgene insertions

Transgene stability following SHS231 integration was analyzed by selection and GFP expression (Fig. 4A). Time-course imaging of GFP fluorescence was performed using an EVOS imaging system (Life Technologies), and the continued expression of SHS231 transgene-encoded Cas9 was quantified by qRT-PCR SYBR green fluorescence on an CFX96 quantitative PCR (qPCR) machine (Cas9 qFwd; 5’-CCCAAGAGGAACAGCGATAAG-3’, Cas9 qRev; 5’-CCACCACCAGCACAGAATAG-3’: BioRad, Hercules, CA). The functional activity of SHS-integrated, transgene-encoded Cas9 protein to promote additional rounds of gene editing was demonstrated by lentiviral transduction and expression of dual gRNAs specific for the *PAX3/FOXO1* fusion oncogene contained in rhabdomyosarcoma cell line Rh30 (Fig. 4B; P/F gRNA1: 5’-GATCAATAGATGCTCCTGA-3’, P/F gRNA2: 5’-GACCTTGTTTTATGTGTACA-3’). The resulting 17.2 kb gDNA-directed deletions were detected using PCR amplification of the region spanning the target gDNA deletion site (Fig. 4B; P/F Fwd: 5’-AGGTTGTCCTGAACGTACCTATCAC-3’ and P/F Rev: 5’-TGCTTCTCCGACACCCCTAATCT-3’; 885 bp).

The functional competence of SHS231 transgene-encoded proteins was further demonstrated using two expression cassettes for the Cas9-based transcription activator proteins dCas9-VPR or Cas9-VPR. Lentiviral expression of dual or triple Cas9 gRNAs was used to target these transactivators to the endogenous, silent *MYF5* gene in Rh5 and SMSCTR cells. The *MYF5* promoter activating gRNAs for dCas9-VPR were gRNA1A, 5’-GATTCCTCACGCCCAGGAT-3’;gRNA2A,5’-GTTTGTCCAGACAGCCCCCG-3’;and gRNA3A,5’-GTTTCACACAAAAGTGACCA-3’. The corresponding truncated activating Cas9-VPR gRNAs targeting the *MYF5* promoter region were tgRNA1A: 5’-GATAGGCTAAAACAA-3’ and tgRNA2A: 5’-GTGCCTGGCCACTG-3’. Changes in *MYF5* gene expression were quantified by SYBR green qRT-PCR using the *MYF5*-specific primers *MYF5* qFwd, 5’-CTGCCCAAGGTGGAGATCCTCA-3’ and *MYF5* qRev, 5’-CAGACAGGACTGTTACATTCGGGC-3’.

The efficiency of SHS231 editing by different endonucleases was determined by co-transfecting two independent RMS cells lines (SMSCTR and RD) with a puromycin-expressing SH231 repair template along with an expression vector for mCreI, for Cas9 nickase (with a single gRNA), or for Cas9 cleavase (with single and dual gRNAs). The RMS cells were also co-transfected with the SHS231 repair template and piggybac transposase plasmid (PB210PA-1, Palo Alto, CA), to compare the SHS231 knockin efficiencies of mCreI and transposase-mediated transgene integration. Two days following transfection, cells were plated into 24 well plates at 3 x 10e4 cells/well, followed by growth in the presence of puromycin (2.5 µg/ml) for 10 days. Cells were then fixed with 2% paraformaldahyde, stained with 0.5% crystal violet and imaged on a Nikon SMZ-745 stereomicroscope to quantify cell number by counting crystal violet stained pixels using imageJ software (NIH).

## RESULTS

### New human safe harbor site identification

Our BLAST search of 128 predicted highly cleavable mCreI target site variants revealed 27 unique mCreI target sites matches in the human genome (Fig. 1A and 1B). A majority of these target sites were found only once (24/27, 89%), while the remaining 3 were represented 2, 3 or 6 times in the human genome for a total of 35 target site matches at different genomic locations (Fig. 1C, Table 2). One of these target sites was a perfect match to a mCreI target site variant (a 20/20 bp match, or 100% identity), whereas the other hits differed by 1 bp (i.e., were 19/20 bp matches or 95% identical) to a query site sequence. The 35 mCreI target sites were located on 16 of the 23 human chromosome pairs including the X chromosome, and covered nearly half of all chromosome arms (23 of 48; Fig. 1C, Table 2).

All 35 new target sites, together with the three canonical human SHS *AAVS1, CCR5* and *hROSA26*, were next evaluated using 8 safety, functional and accessibility criteria in addition to site uniqueness (Table 1 and 2). Among our 35 newly identified sites, 25 (or 71%) fulfilled more than half (≥5/9) of our SHS criteria, as did the *AAVS1* and *CCR5* canonical human SHS (Table 2). When we examined safety criteria alone (SHS criteria 1-6 in Table 1), 21/35 (60%) of our target sites met ≥4 of 6 criteria, with three (SHS231, 233 and 303) matching all 6 safety criteria. In contrast, the widely used human SHS *AAVS1, CCR5* and *hROSA26* each matched only 3 of 6 safety criteria (Table 2). This site assessment was more extensive than previous attempts and made systematic use of genomic data that together, allowed us to rank-order both newly identified and canonical SHS for potential utility and experimental verifications (Table 2).

Genetic variation between individuals has the potential to complicate or disrupt the editing of SHS as well as other genomic regions. In order to assess the potential magnitude of this problem, we assessed all 35 of our new SHS for copy number and basepair-level genetic variation. None of our target sites was located in a copy number-variable region of the human genome, though we did identify base pair-level genetic variation in 10 of our 35 mCreI target sites in whole genome sequencing data generated as part of the 1000 Genomes Project^21^. This site-specific base-pair variation was restricted to single nucleotide polymorphic variants (SNPs or SNVs); no indels were identified. Four SHS contained potential mCreI cleavage-inactivating SNP variants: SHS255 on chromosome 5 (variant frequency = 0.5041), SHS301 on chromosome 7 (variant frequency = 0.2234), SHS293 on chromosome 8 (variant frequency = 0.0037) and SHS297 on chromosome 17 (variant frequency = 0.0751). All four SNPs were predicted to strongly suppress mCreI cleavage efficiency by ≥70% (Fig. 1B, Table S2). Of note, among individuals analyzed as part of the 1KGP, 80% lacked any SNP variants in *any* of our 35 target sites including SHS231, and 94% had all 35 target sites predicted fully mCreI-cleavage sensitive despite the presence of one or more permissive base-pair variant SNP (Table S2 and additional results not shown).

### Experimental validation of potential new human SHS

In order to experimentally validate the most promising of our potential new SHS, we amplified 28 of the target site regions from the human genome and subjected these to either *in vitro* mCreI cleavage assays or DNA sequencing. As part of these analyses we identified one polymorphic 108 bp insertion adjacent to SHS231 that was present in a subset of human cell lines. This insertion contained a 35-base poly-T sequence and adjacent short sequence blocks reminiscent of transposable element short tandem duplications, and was found to be an exact match for a segment of an AluYa5 subfamily, SINE-derived repeat of 311 bp that is present in ∼4000 non-redundant copies in the human genome (see: http://www.dfam.org/entry/DF0000053). Though located near SHS231, we demonstrate below that this insertion did not affect SHS231 access or editability. A majority of SHS were fully cleavage-sensitive *in vitro* when compared with the canonical mCreI target site, including single copy SHSs 227, 229, 231, 233, 251, and multi-copy SHSs 253, 255, 257, 259, 263. As noted above, all of the individuals analyzed as part of the 1KGP either lacked *any* SHS SNP variants (80%), and 94% had all 35 sites predicted fully mCreI-cleavage sensitive (Table S2; additional results not shown).

### Efficient *in vivo* cleavage and editing of new SHS by multiple genome editing nucleases

We assessed the functional competence of potential new SHS by determining their *in vivo* cleavage sensitivity and ability to be edited by different genome editing nuclease/repair template combinations. These experiments focused on the single copy, highly-ranked chromosome 4q SHS231, and two sites on chromosome 2 that were single copy (SHS229), or as a single copy on chromosome 2 with additional copies on chromosome arms 5p, 7q, 14q, 17q and Xp (SHS253; Fig.1, Table 2). The *in vivo* cleavage sensitivity of these and three additional SHS was analyzed by co-expressing mCreI with the TREX2 3′ to 5′ repair exonuclease in human 293T cells, followed by PCR amplification and mCreI digestion of target sites. This experiment was designed to identify a cleavage-resistant target site fraction in nuclease-expressing cells, from which a minimum estimate of *in vivo* cleavage efficiency can be derived^22^.

Five of the 6 SHS assayed in this way, the unique sites SHS227, 229 and 231 and copies of the same target site sequence located on different chromosomes (SHS253, 257 and 263), had increased fractions of mCreI-resistant target site PCR products that ranged from 3.8% to 31.3% when compared with the corresponding SHS-specific PCR product from mock-transfected control cells. The presence of multiple SHS-specific, mCreI-resistant PCR products also provides evidence for the ability of mCreI to cleave-and thus potentially simultaneously edit-multiple target sites in human cells.

In order to determine whether SHS cleavage *in vivo* could catalyze high fidelity homology-dependent repair, we co-transfected human 293T cells with an expression vector for mCreI, for a CRISPR/Cas9 cleavase/nickase or for a TAL effector nuclease (TALEN) pair together with a SHS-specific repair template containing a *loxP* site flanked by two different diagnostic restriction sites (Fig. 2). SHS229, 231 and 253 were analyzed following mCreI expression, SHS229 and 231 after CRISPR/Cas9 cleavase/nickase expression, and SHS231 after TALEN expression. PCR amplicons from transfected cells were then subjected to PvuI and SacII restriction digestion to confirm targeted capture and site-specific integration of the *loxP* repair template, followed by cloning and DNA sequencing to confirm the structure and fidelity of cleavage-dependent, targeted SHS integration (Fig. 2). The frequency of targeted SHS231 integration events in 293T cells was 4.8% for mCreI/TREX2 (3/63 clones); 6.1% (2/33) for CRISPR/Cas9 nuclease and 16.1% (5/31) for CRISPR/Cas9 nickase; and 1.23% (1/81) for a SHS231-specific TALEN pair (Fig. 2). Infrequent single base substitutions observed in cloned and sequenced *loxP* inserts were most likely PCR errors introduced by Taq DNA polymerase during site amplifications for cloning and DNA sequencing. Parallel targeted integration assays at SHS229 and 253 showed comparable results (Fig. S2).

In order to increase SHS engineering efficiency and potentially facilitate the editing in post mitotic cells, we also evaluated SHS231 editing by a potentially homology-independent knockin approach. This strategy used Cas9-mediated cleavage of the repair template and genomic SHS target locus (i.e., using dual gRNAs; US2-Cas9) to promote potential repair with transgene integration by NHEJ-mediated repair mechanisms^32,33^ (Fig. 3A). While indel mutations can be introduced during NHEJ-mediated repair in the cleaved target locus and repair template, this is not a serious concern since our SHS were specifically identified to contain no functional genomic elements and the repair template cleavage site did not inactivate the encoded transgene(s). Molecular analysis of SHS231 integration events by amplification, cloning and sequencing of the 5’ SHS231 integration site identified both direct fusion events (no indels), as well as the expected short indel mutations at the gRNA cleavage site (Fig. 3A), evidence compatible with NHEJ-mediated integration. The efficiency of dual gRNA Cas9 cleavage-mediated editing of the SHS231 locus was compared to the Cas9 nickase, cleavase and mCreI-mediated HDR approaches by co-transfection of each endonuclease with a repair template expressing puromycin (Fig. 3B-C, Fig. S1). The efficiencies of these endonucleases was also compared to random integration of the repair template using a piggybac transposon, since the repair template contained piggybac terminal repeat sequences flanking the transgene cassette. This experiment was performed in two independent RMS cells lines (RD and SMSCTR), where the putative homology-independent insertion or knockin of the puromycin repair template was 2-fold higher when compared to HDR-mediated insertion. Neither of these approaches, however, was as efficient as random integration by piggybac-mediated transposition (Fig. 3B and 3C).

**Figure 3.**
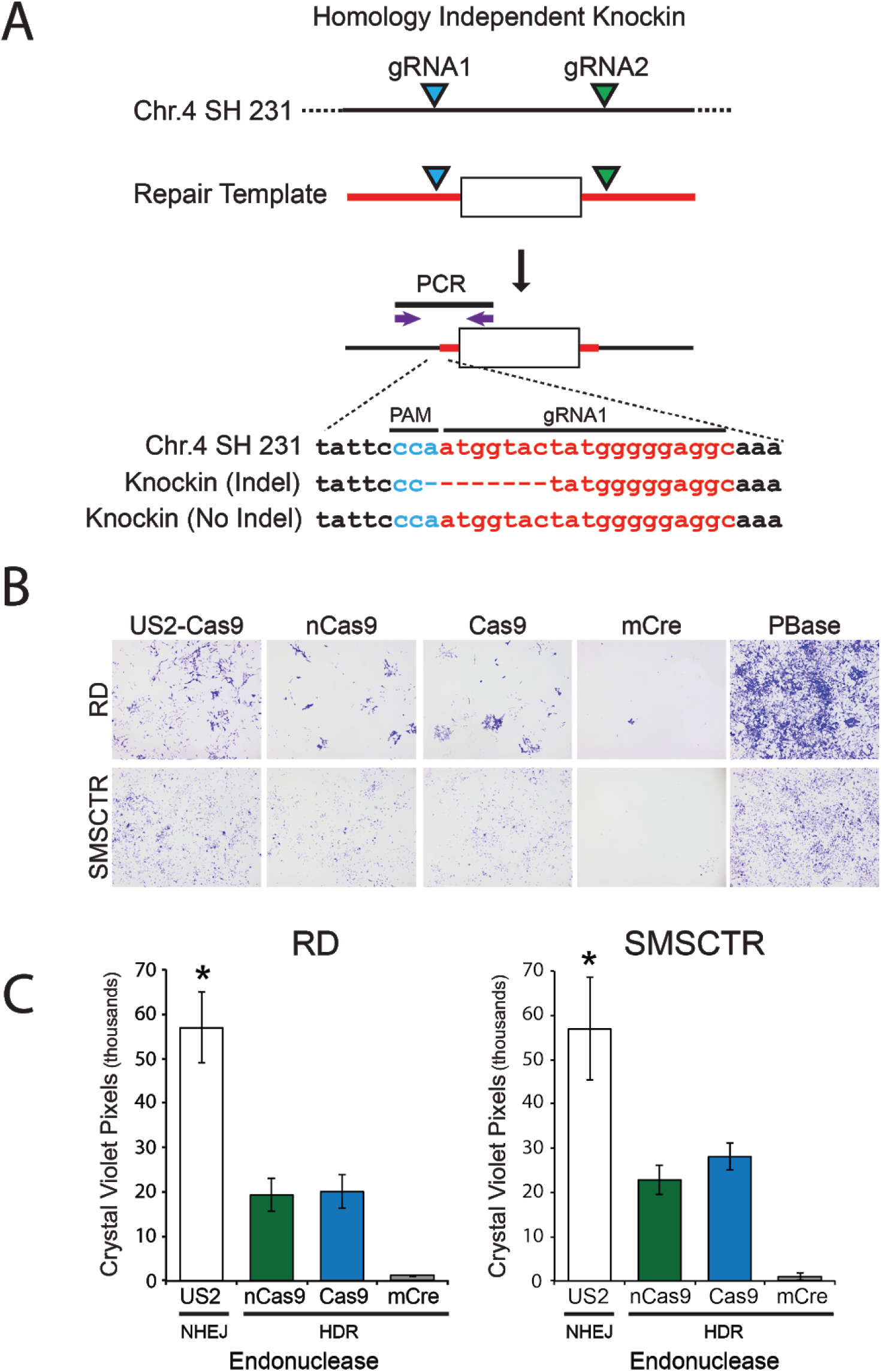
Homology-independent engineering of the chromosome 4q SHS231. **(A)** Strategy for targeted integration of transgene cassettes using NHEJ mediated repair. Blue and green triangles represent gRNA target sites on both the genome and repair template. Representative sequences from the 5’ transgene integration site after knockin specific PCR amplification of an integrated transgene (purple arrows). **(B)** Relative knockin efficiency of a puromycin cassette using homology independent repair (US2-Cas9; NHEJ), and homology directed repair (nCas9, Cas9, mCreI; HDR) at the SHS231 locus, compared to piggybac transposition (PBase). **(C)** Quantification of crystal violet staining from SHS231 knockin stable cells. * Significantly different from HDR SHS231 knockin approaches, P<0.05.

### Characterization of stability, expression, and functionality of SHS231 integrated genes

The functional utility of any SHS depends critically upon persistent marking and/or SHS-specific gene expression after site editing. In order to assess this key SHS functional requirement, we analyzed the expression of several different transgene cassettes that had been integrated into the chromosome 4 SHS231. SHS transgene expression stability was assessed by integrating, and then following the expression of, a SHS231 GFP reporter cassette in two independent RMS cells lines (SMSCTR and Rh5) where transgene insertion was mediated by putative homology-independent editing. When GFP transgene expression was followed over several weeks (i.e., over 45 days) in the absence of antibiotic selection, we observed no significant decrease in GFP expression after 15 population doublings (Rh5) or 25 population doublings (SMSCTR; Fig. 4A). These results highlight the stable nature of transgene integration and expression from SHS231, over usefully long periods of time in mitotically dividing cells.

**Figure 4:**
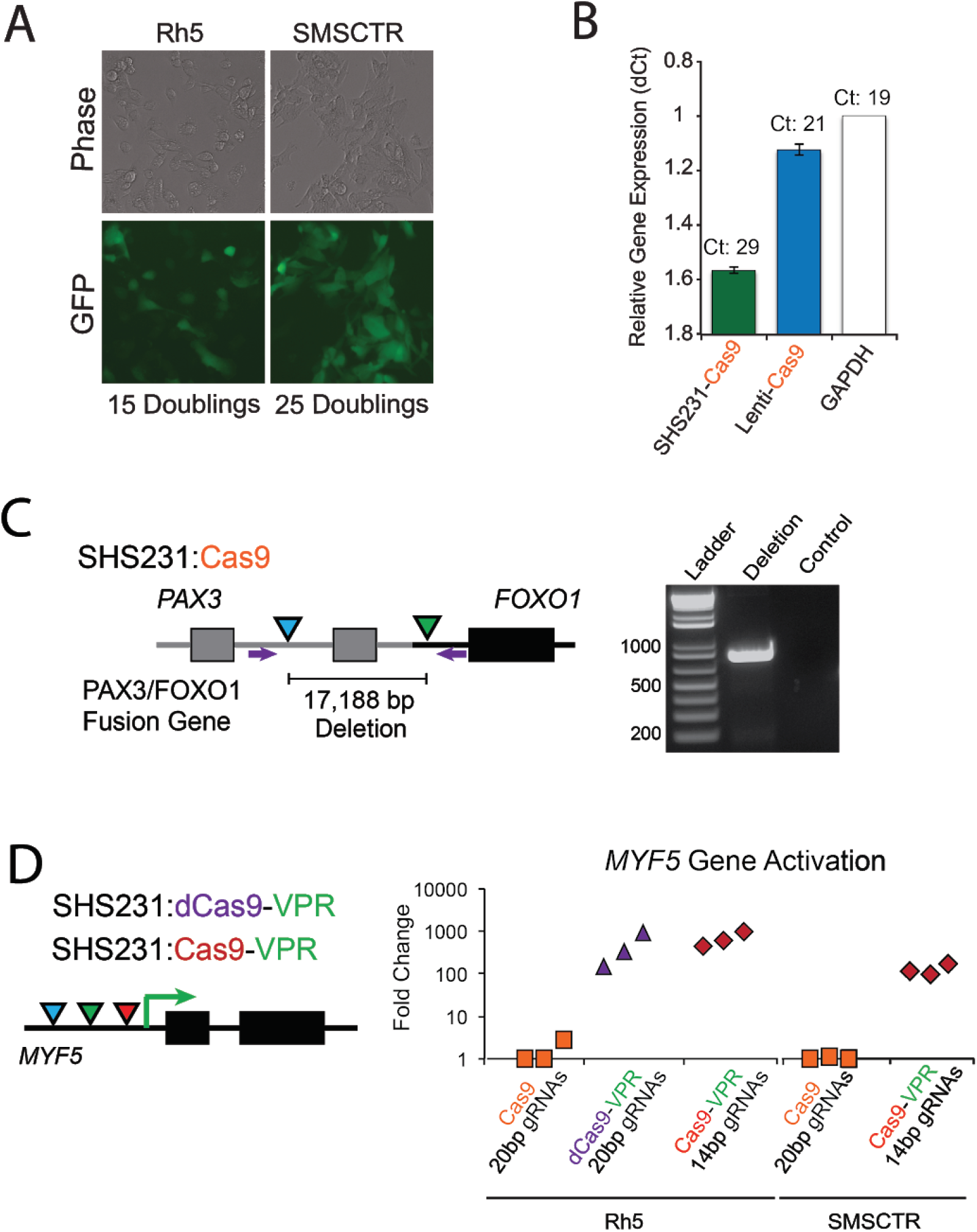
Stable expression of functional gene editing and gene activation proteins encoded by SHS231 transgenes. **(A)** Long-term stable GFP expression from a SHS231 integrated transgene in two independent RMS cell lines. **(B)** Relative Cas9 expression level (cycle threshold: Ct) from a SHS231 integrated Cas9 cassette compared to cells transduced with high titer Cas9 expressing lentivirus or the endogenous expression level of GAPDH. Both SHS231 and lentiviral Cas9 variants were expressed from the human EF1a promoter. **(C)** Targeted deletion of a 17,188 bp gDNA segment of the PAX3/FOXO1 fusion oncogene in Rh30 RMS cells expressing Cas9 from the SHS231 locus. Dual gRNA target sites (blue and green triangles) and deletion PCR primer sites (purple arrows) are identified. **(D)** Demonstration of endogenous *MYF5* gene activation with SHS231 expressed dCas9-VPR and Cas9-VPR transgenes. Gene activation was achieved by targeting full length (20bp) or truncated (14bp) gRNAs (blue, green, and red triangles) to the promoter region of the MYF5 gene.

We next determined whether SHS231-integrated, Cas9-derived transgenes were not only persistently expressed but retained their intended functions. Stable Cas9-expressing cell lines are a convenient starting point for a growing range of Cas9-enabled methods to study gene structure, function or to enable genetic screens. We observed readily detectable Cas9 expression from SHS231 knockin transgenes that was comparable to cells super-infected with high titer lentivirus to express Cas9 protein, or to the expression of endogenous GAPDH protein (Fig. 4B). The functional competence of SHS231-expressed Cas9 protein was further demonstrated in Rh30 RMS cells by transducing cells with a lentivirus expressing two gRNAs targeting a *PAX3/FOXO1* fusion oncogene contained in Rh30 (Fig. 4C). Efficient generation of the predicted 17,188 bp gDNA-targeted deletion in *PAX3/FOXO1* was readily detected by PCR amplification of gRNA-transduced cell pools using primers that flanked the *PAX3/FOXO1* gRNA target sites (Fig. 4C).

In a third series of SHS functional validation experiments, we integrated transgene cassettes in SHS231 that expressed chimeric Cas9-derived transcriptional activators dCas9-VPR or Cas9-VPR by Cas9-mediated knockin. VPR is a tripartite transcription factor consisting of VP64, P65 and Rta transactivation domains^34^. Fusion of this transcription factor to the C-terminus of the Cas9 protein generates a potent, programmable transcriptional activator (dCas9-VPR or Cas9-VPR)^34^. Each SHS231 RMS cell line expressing dCas9-VPR or Cas9-VPR was then transduced with a lentivirus expressing 2 or 3 gRNAs targeting the promoter region of the *MYF5* gene (Fig. 4D). *MYF5* is typically not expressed or expressed at very low levels in many RMS cells, and therefore is a good candidate for measuring gRNA-targeted Cas9-VPR-mediated gene activation. We found that both full length (20bp) and truncated (14 bp) gRNAs promoted robust Cas9-VPR-dependent *MYF5* gene activation in both of the RMS cell lines tested (Fig. 4D).

These results collectively demonstrate efficient editing of a newly defined human safe harbor site, and the stable expression of functionally useful SHS231-integrated transgenes encoding GFP and Cas9 protein variants. Moreover, we demonstrate the ability of these proteins to drive additional useful outcomes including genome editing with the promotion of large deletions in a *PAX3/FOXO1* fusion oncogene, and induced expression of the *MYF5* gene that is normally silent in RMS cells. The SHS231-specific targeting vectors used in these experiments have been assembled into a SHS231-specific ‘toolkit’ to enable facile editing of the highly-ranked SHS231 in a wide range of human cell types (Fig. S1, Table S3). This SHS231 toolkit is available from Addgene (Addgene, Cambridge, MA), and includes both Cas9 and dCas9-based expression cassettes, as well as GFP and RFP reporter constructs with puromycin, hygromycin and blasticidin selectable markers. All of the expression vector transgenes included in this set are driven by the human EF-1α promoter and contain additional *attP* sites to serve as ‘landing pads’ for ΦC31 and Bxb1-mediated, high efficiency SHS transgene insertion.

## Discussion

Only a small number of SHS are in wide use in human cells. These were originally identified by serendipity (*AAVS1, CCR5)* or by their similarity to SHS in other organisms (e.g., *hROSA26)*. In order to address the continuing need for additional well-validated human SHS to enable a broader range of basic and translational science applications, we used a systematic approach to identify and evaluate 35 potential new SHS in the human genome. These new SHS cover a substantial fraction of the human genome: 16 of 23 chromosomes including the X chromosome, with SHS on 23 of 48 chromosome arms (Figure 1). These potential new SHS were assessed and rank-ordered as potential ‘safe harbors’ using both previously suggested criteria (e.g.,^17^) and additional more recently available human genome-scale structural, genetic and regulatory data (e.g., ENCODE data^18^). Over half of our new SHS (20/35, or 57%) met 4 of our 6 core safety criteria (Tables 1 and 2), in contrast to the widely used human *AAVS1, CCR5* and *hROSA26* SHS that each met 3 or fewer of these core safety criteria (Table 2).

All 35 of these newly identified SHS contained a site-anchoring 20 bp mCreI nuclease cleavage site, and thus can be immediately targeted either singly or in multiplexed fashion using this small, easily vectorized homing endonuclease together with SHS-specific repair templates^7–9^. All of these SHS can also be targeted by virtue of overlapping or adjacent Cas9 and TALEN target sites, as we demonstrated for three different sites located on chromosomes 2 and 4. Of note, human population genomic data indicate that few of these 35 new human SHS harbor any genetic variation that would prevent their use for mCreI, Cas9 or TALEN-mediated editing in human cells or cell lines.

As part of the experimental validation of a subset of these new human SHS, we demonstrated both Cas9 nickase and cleavase-dependent editing, and efficient editing of the chromosome 4 SHS231 by both homology-dependent and likely homology-independent, NHEJ-mediated mechanisms. High efficiency, homology-independent transgene integration strategies in which both template and target locus are cleaved may facilitate higher efficiency site-specific editing while taking advantage of the less stringent requirements for editing than endogenous open reading frame editing by higher fidelity homology-dependent approaches. Thus a dual-cleavage knockin approach may facilitate the efficient generation of cell populations with virtually identical, site-specific transgene insertions. This approach could in many instances eliminate the time and expense of isolating multiple cell clones, while retaining the natural heterogeneity found in the human cells and cell lines most often used to study and model biological systems. Dual-cleavage knockin strategies also have the potential to open many non-dividing cell types to efficient genome engineering, in contrast to homology-dependent pathways that can only be efficiently used in dividing cells.

Several aspects of our newly defined SHS remain to be explored and/or optimized. While we have thus far extensively validated only a subset of our sites (SHS231, 229 and 253; Figure 1), we anticipate these sites will be representative of most or all of our other newly identified SHS in different cell types. Most notable among these results was targeted transgene insertion with persistent expression from SHS231 of useful transgene-encoded proteins such as Cas9 variants, selectable markers and fluorescent proteins. Stable transgene expression is a key requirement for SHS, and thus will need to be further verified to identify SHS-specific variables that might affect SHS editing and transgene expression in different cell types (see, e.g., Daboussi *et al.,* 2012^38^). Should site-specific problems arise, the substantial expansion of useful new human SHS identified here may provide ready experimental alternatives.

The efficiency of SHS-targeted editing can likely also be further optimized. Important variables include cell type-specific gene transfer efficiencies; repair template type (single-vs double-stranded), and the length and degree of nucleotide sequence identity between the repair template and target site flanking sequences. The highest efficiency of homology-directed repair can in most instances be promoted by incorporating >200bp of perfect DNA sequence identity between a SHS and donor repair template arms^39–42^. Thus target site characterization in cell types of interest is an important part of any homology-dependent editing optimization workflow, in order to identify potentially confounding issues such as the variable SINE/Alu-derived short insertion we identified near the SHS231 site in a subset of cell lines. This type of unanticipated finding, once identified, can be readily incorporated into the construction of repair templates where long, flanking homology arms are desirable or required.

The new SHS identified here expand by an order of magnitude the number of human SHS that can be used for human genome editing and engineering applications. The SHS assessment and scoring strategy we used was more comprehensive that previous efforts, and can be further modified to incorporate new or application-specific SHS scoring criteria. For example, the growing number of apparently dispensable human genes^6,43^ offers one rich source of potential new human SHS. These human gene ‘knockout’ lists can be supplemented with complementary lists of essential or high fitness human genes, to focus on genomic regions to target or avoid as part of genome engineering projects^44-46^. The characterization of additional new human SHS and the development of SHS-specific reagents such as our SHS231 ‘toolbox’ should provide practically useful tools to enable a wide range of basic as well as translational human genome engineering applications.

## Acknowledgements

We thank Drs. Kenny Matreyek and Douglas Fowler for introducing us to the BxbI recombinase system and for providing materials; Alden Hackmann for help with Figures; and Marilyn Moelhman and Texia Loh for work on the design and analysis of CRISPR activation and SHS231 homology independent knockin experiments. SP, HL, RBS and RJM Jr were supported by NIH Award 1RL1CA133831, while BTH was supported by Interdisciplinary Training in Genomic Sciences NHGRI T32 Award HG00035. MP and EC were supported by NIH R01 CA227432-01.

## Author Disclosure Statement

None of the other authors has a competing interest to declare.

---

**Figure S1.**
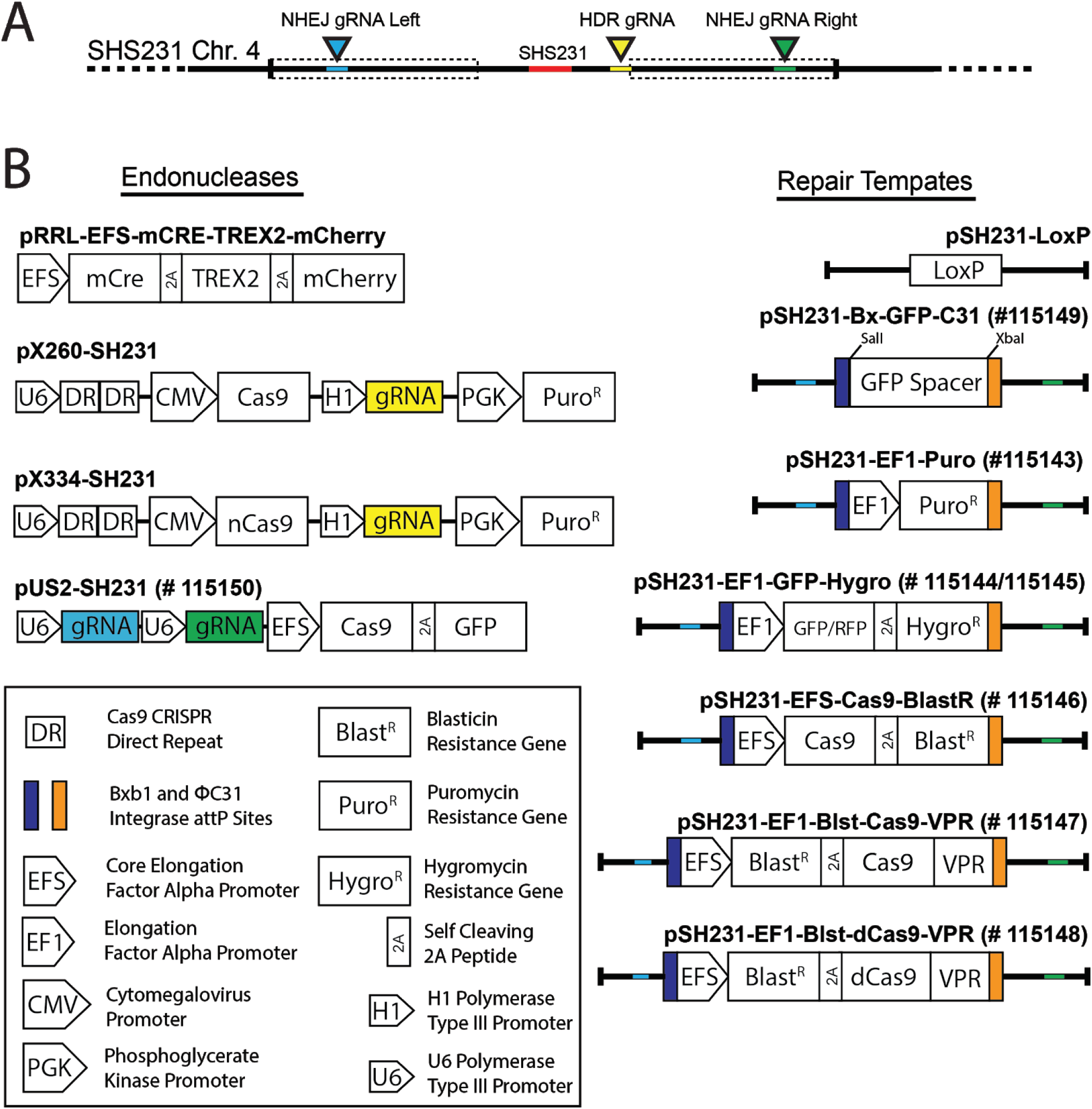
SHS231 endonuclease and repair template constructs. **(A)** Details of the SHS231 locus with homology dependent (HDR) and homology independent (NHEJ) gRNA target sites identified along with the location of repair template homology arms (dashed boxes). **(B)** Features of the endonuclease expression and repair template vectors are identified in the legend. The gRNA colors correspond to target sites in the safe harbor locus and in repair template homology arms.

**Figure S2:**
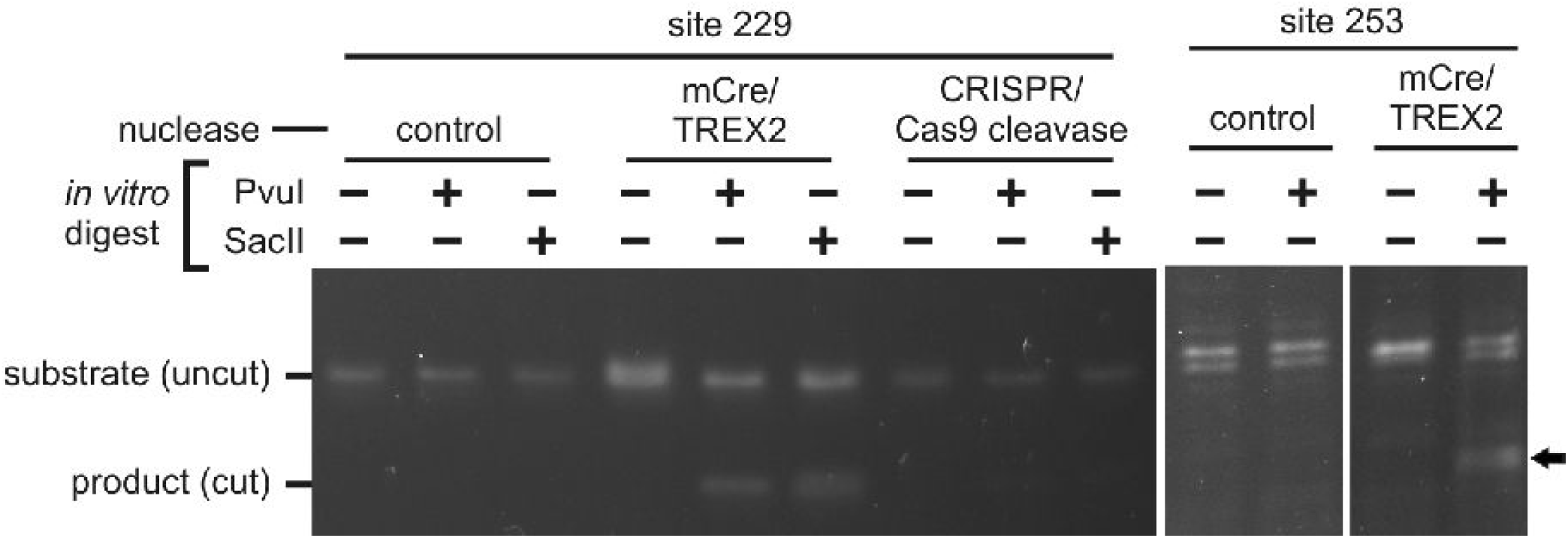
Restriction site analysis from HDR integration of a loxP cassette into the SHS229 and SHS253 loci.

**Table S1.**
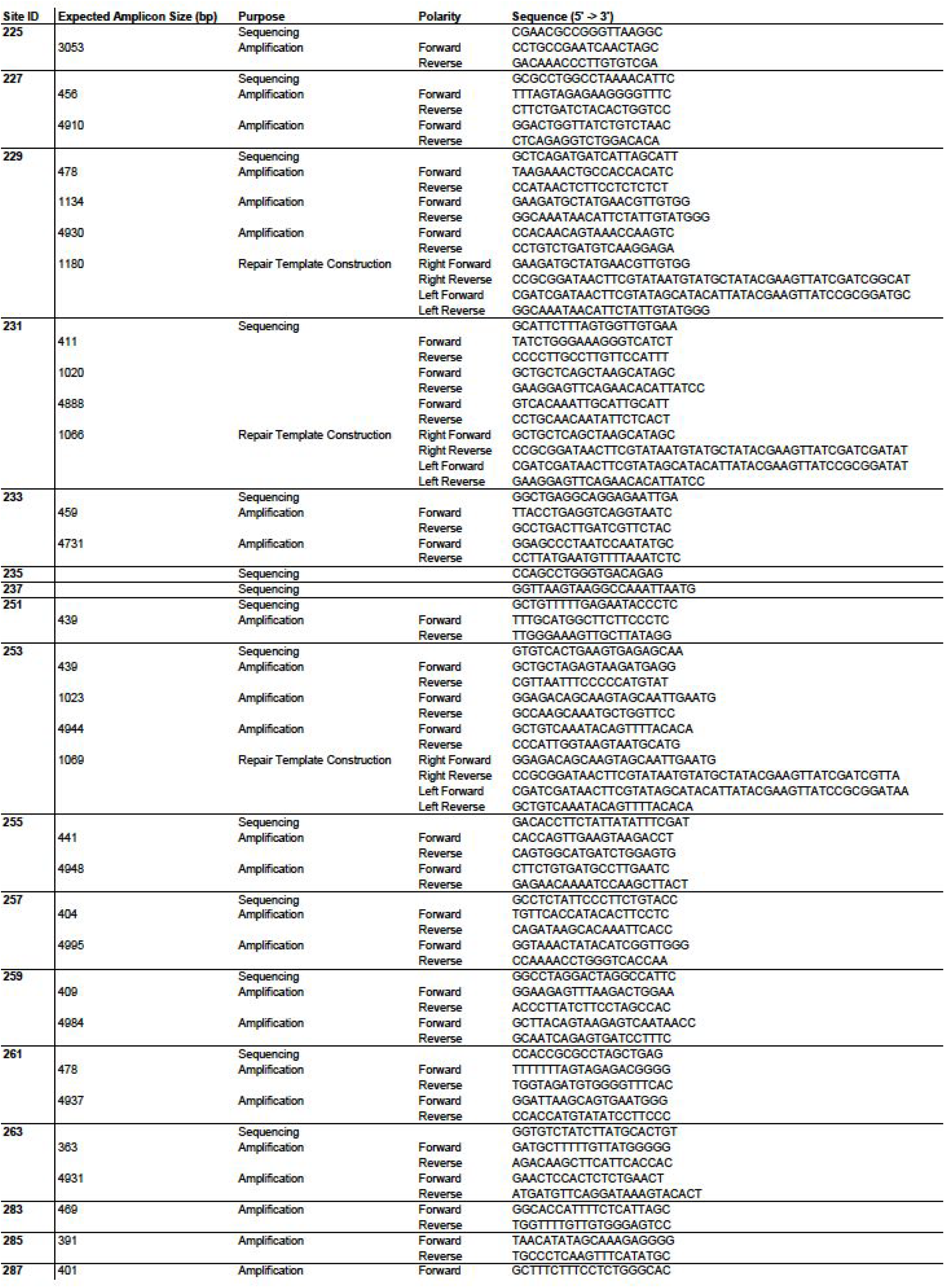
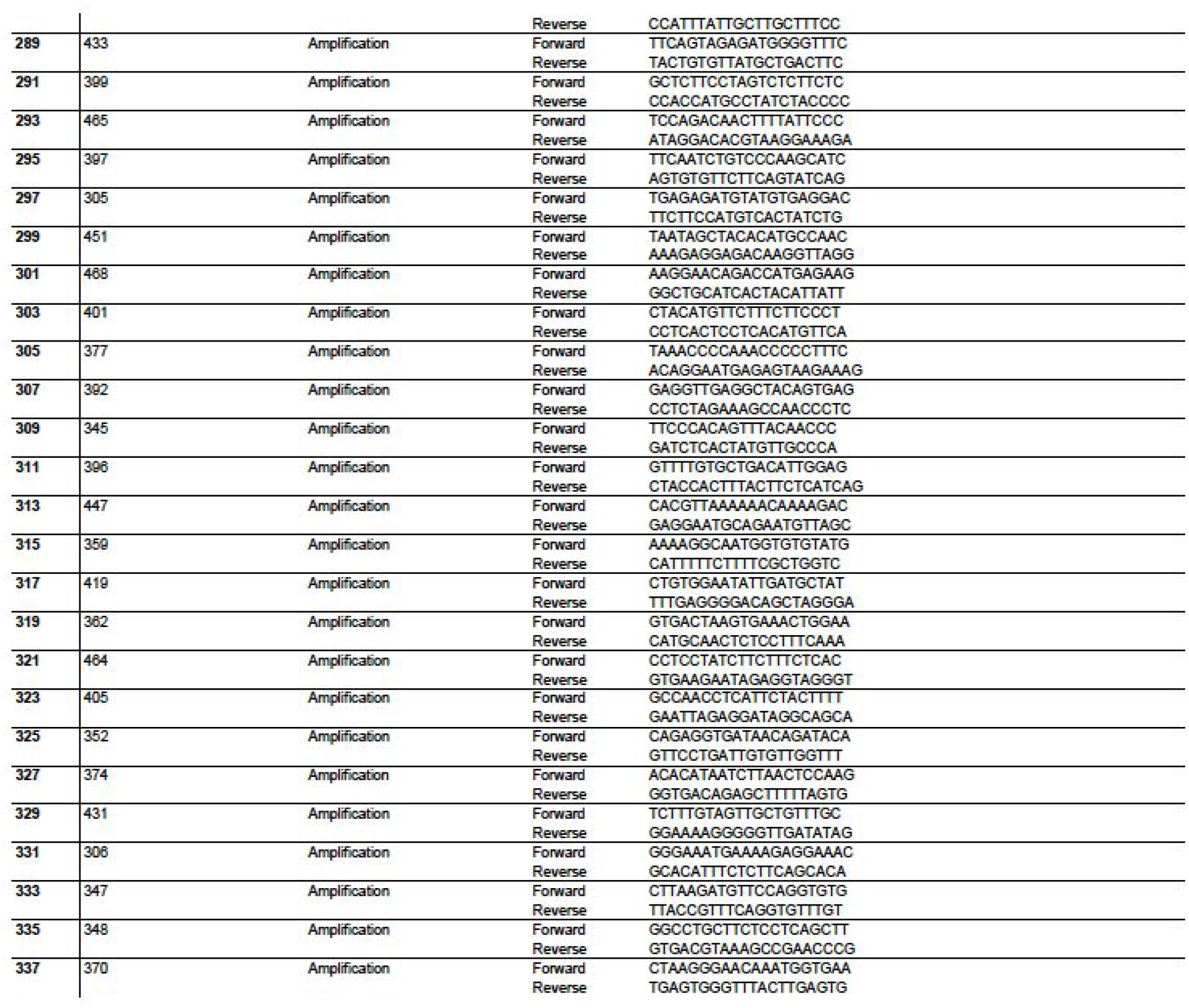
Sequences of primers used for SHS amplification, sequencing, and vector construction.

**Table S2:** Nucleotide sequence variants in mCreI genomic target sites together with predicted effect on mCreI cleavage sensitivity.

| Site ID | Chr | Start | End | SNV Position | SNP | Cre position | Frequency | Effect |
| --- | --- | --- | --- | --- | --- | --- | --- | --- |
| 323 | 1 | 152360840 | 152360859 | 152360844 | C/T | G @ + 6 (rev) | 0.000457875 | 0.81 |
| 229 | 2 | 45708354 | 45708373 | 45708365 | C/T | C @ + 2 | 0.002289377 | 0.99 |
| 283 | 4 | 37769238 | 37769257 | 37769243 | A/G | A @ -5 | 0.000457875 | 0.69 |
|  |  |  |  | 37769246 | A/G | A @ -2 | 0.000457875 | 1.21 |
| 315 | 5 | 7577728 | 7577747 | 7577738 | A/G | C @ -1 (rev) | 0.007326007 | 0.59 |
| 255 | 5 | 19069307 | 19069326 | 19069307 | A/G | G @ -10 | 0.504120879 | 0.28 |
| 305 | 5 | 159922029 | 159922048 | 159922040 | C/T | G @ -2 (rev) | 0.009157509 | 1.00 |
| 301 | 7 | 113327685 | 113327704 | 113327699 | C/T | T @ 5 | 0.223443223 | 0.21 |
| 257 | 7 | 138809594 | 138809613 | 138809604 | A/G | C @ -1 (rev) | 0.000457875 | 0.59 |
| 293 | 8 | 40727927 | 40727946 | 40727939 | A/G | T @ -3 (rev) | 0.003663004 | 0.17 |
| 297 | 17 | 14810285 | 14810304 | 14810291 | C/T | C @ -4 | 0.075091575 | 0.16 |

**Table S3:**
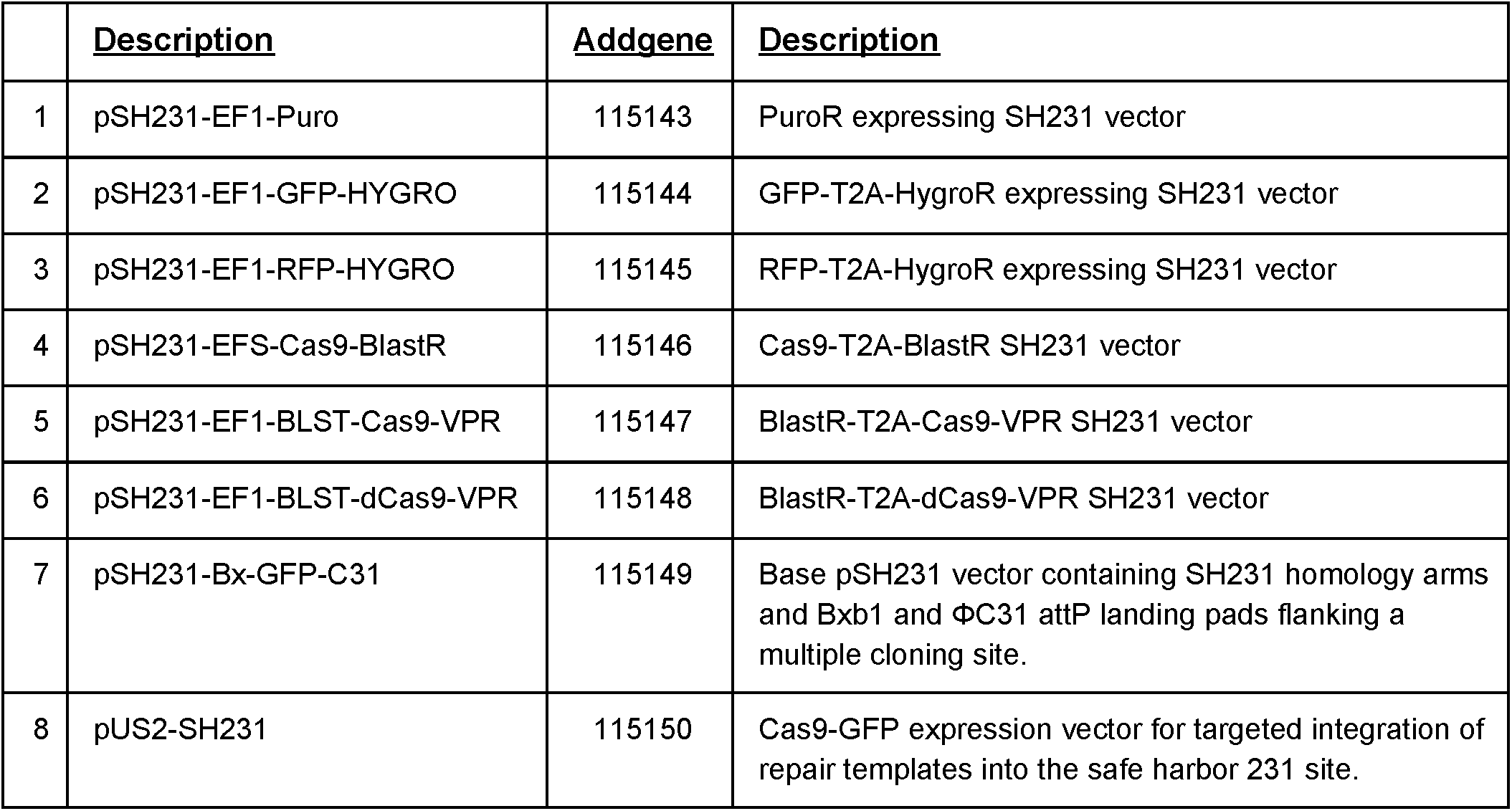
Human chromosome 4 SHS231 editing toolkit.

